# Using expert-elicitation to deliver biodiversity monitoring priorities on a Mediterranean island

**DOI:** 10.1101/2021.08.18.456866

**Authors:** J. Peyton, M. Hadjistylli, I. Tziortzis, E. Erotokritou, M. Demetriou, Y. Samuel-Rhoads, V. Anastasi, G. Fyttis, L. Hadjioannou, C. Ieronymidou, N. Kassinis, P. Kleitou, D. Kletou, A. Mandoulaki, N. Michailidis, A. Papatheodoulou, G. Payiattas, D. Sparrow, R. Sparrow, K. Turvey, E. Tzirkalli, A.I. Varnava, O.L. Pescott

## Abstract

Biodiversity monitoring plays an essential role in tracking changes in ecosystems, species distributions and abundances across the globe. Data collected through both structured and unstructured biodiversity recording can inform conservation measures designed to reduce, prevent, and reverse declines in valued biodiversity of many types. However, resources for biodiversity monitoring are limited, it is therefore important that funding bodies prioritise actions relative to the requirements in any given region. We addressed this prioritisation requirement through a three-stage process of expert-elicitation, resulting in a prioritised list of twenty biodiversity monitoring needs for Cyprus. Equal priority was assigned to the twenty monitoring needs within three categories: a top nine, a middle five, and a bottom six. The most highly prioritised biodiversity monitoring needs were those related to the development of robust methodologies, and those ensuring a geographic spread of sufficiently skilled and informed contributors. We suggest ways that the results of our expert-elicitation process could be used to support current and future biodiversity monitoring in Cyprus.

## Introduction

The earth’s climate and habitats are changing at unprecedented rates, with species and ecosystems increasingly threatened by multiple, often interacting, anthropogenic pressures (1). Five main threats to biodiversity were identified by the Intergovernmental Science-Policy Platform on Biodiversity and Ecosystem Services (IPBES): changes in land and sea use; direct exploitation of organisms; climate change; pollution; and, invasive alien species (IAS) (1).

The Mediterranean basin is one of the 34 global biodiversity hotspots, including a large number of endemic species where, due to increased anthropogenic pressures, there is an urgent need for species and habitats conservation (2). Mediterranean-climate areas are predicted to be likely to experience the highest levels of biodiversity change by 2100, due to pressures such as changes in land use (e.g. agriculture), climate change, and the introduction of IAS (3). The most conspicuous threats that affect the greatest number of taxonomic groups in the Mediterranean are habitat loss and degradation, which are considered the primary impacts on diversity, followed by exploitation, pollution, climate change, eutrophication and species invasions (4). Increased tourism and urbanisation in this region also adversely affect biodiversity through habitat loss, disturbance, use of pesticides and herbicides and increased pollution (5–7). Over-exploitation of natural resources is another example of the effects of anthropogenic pressure on species in the Mediterranean including fisheries of the Mediterranean Sea, which are at risk of over-exploitation, with a pressing need for assessments of stock numbers (8).

To attempt to halt and, where possible, reverse species population declines and habitat degradation, policy-makers need to use scientifically robust biodiversity data to understand the current situation (9, 10). Monitoring is therefore needed to provide evidence on biodiversity status and trends (11). Hochkirch, Samways (12) suggest an eight-point strategy for reducing gaps in such biodiversity data, of which the most relevant at the local level are: increasing explorative field surveys; increasing monitoring of less well-studied taxa; building capacity in areas where there is high species-richness or levels of endemism; and the use of global online data repositories such as the Global Biodiversity Information Facility (GBIF; https://www.gbif.org/) for hosting high quality, open data. Open data can be an important mechanism enabling the effective use of biological monitoring data (13). The increasing use of technologies for biological recording has resulted in many millions of records being made freely available through global platforms such as GBIF. However, open data repositories may also contain important biases in the records uploaded (12, 14).

Information on species distributions and identification (15) is essential for supporting conservation (16), but funding and support for taxonomic training may be limited (17) with numbers of taxonomists decreasing globally (18, 19). Technological methods are being increasingly used to fill knowledge gaps, as they may assist with the surveying of large or difficult to reach areas more rapidly and cheaply than on-the-ground surveys, e.g. through remote sensing (20), the use of habitat suitability maps (2), and environmental DNA (21). Initiatives such as the Distributed European School of Taxonomy, which offers education and training opportunities (https://cetaf.org/dest/about-dest/), are another mechanism for bridging these gaps; however, limited support for both taxonomic training and the availability of employment still present challenges for conservation (22, 23). Indeed, the “provision of funding mechanisms” was another strategy highlighted by Hochkirch, Samways (12) as important for increasing the availability of biodiversity data.

Reduced governmental funding for the collection and curation of records within statutory organisations in the UK has led to a diversification in mechanisms of biological data gathering (24). Citizen science, the involvement of volunteers in generating scientific data (including biological records) is often seen as an attractive option by policy-makers to fill knowledge gaps in biodiversity datasets (25). Citizen science can provide broad geographic coverage of species’ occurrences (26, 27) and has been used to monitor trends in the distributions (28, 29) and abundances of species (30–32). The provision of early warnings of the arrival of new IAS is another area of success for citizen science (33–35). Even in regions without long-standing traditions of amateur contributions to natural history, important discoveries are still being made in this way. For example, a new marine alien species to Cyprus was discovered by recreational divers in 2019, and was identified through photographs posted in the online data repository of the iSea project “Is it Alien to you? Share it!!!”(36).

As noted above, despite the existence of open databases on species distributions, new technologies, and the diversification of mechanisms for collecting monitoring data, knowledge gaps still exist (37, 38). Where existing data and models cannot answer questions needed to inform management and policy, expert-elicitation, i.e. using the knowledge and experience of experts, can be used to address these gaps (39). However, the use of expert judgement can bring its own biases into such decision-making processes (40). Using structured approaches to expert-elicitation may help to overcome these biases; Roy, Peyton (41) outline ten guiding principles to reduce bias during expert-elicitation processes. Expert-elicitation methods, such as the Delphi technique, have been used to address many questions in conservation science (42), including the prioritisation of knowledge needs for enabling implementation of “nature-based solutions” in the Mediterranean (43). Expert-elicitation methods have also been used as a mechanism for prioritising needs in biodiversity monitoring (44). Prioritising biodiversity monitoring needs for data contributors (whether amateur or professional) and end users of data (e.g. governments) can prove a useful mechanism for mediating between potentially competing priorities between such groups (44), and can help to focus spending where resources are limited.

Here we sought to use expert-elicitation approaches to collaboratively prioritise the biodiversity monitoring needs of a range of stakeholders on a Mediterranean island, rich in biodiversity. This was undertaken both in order to increase understanding of biodiversity monitoring options that could be utilised, and to provide a stable list of priorities on which to base future policy development, resource allocation, and general decision-making in this area.

## Study area

Cyprus is situated 65 km south of Turkey and 105 km west of Syria; it covers an area of 9,251 km^2^ (45). Cyprus is very rich in biodiversity in relation to the size of the island (46), with a variety of landscapes, species and habitats of European importance, and has high cross-taxon levels of endemism (47–50). Plants in particular have one of the highest levels of endemism in the European Union (51, 52).

The Government of the Republic of Cyprus and Sovereign Base Area Administration fund, among other initiatives, the monitoring of protected species and habitats on Cyprus. However, to date, as throughout the world, the biodiversity recording effort on Cyprus has been uneven, with many geographical areas and taxonomic groups poorly represented on biodiversity data platforms, particularly at finer spatial scales. Whilst structured biodiversity monitoring does exist on the island, this is generally aimed at generating temporal trends in population counts, and is largely restricted to birds, although we also note the monitoring activities of the Game and Fauna Service of the Republic of Cyprus Government of the endemic mouflon going back for over two decades. BirdLife Cyprus has been publishing annual reports of unstructured bird records since 1970, as well as monthly checklists based on reports by birdwatchers. Their longest running structured monitoring scheme is the monthly water bird survey, which started in 2005. The Cyprus Dragonfly Study Group and the Cyprus Butterfly Group have also recently begun to contribute data to relevant pan-European projects (53, 54). Plants have been well-recorded at a broad scale (55), but, again, finer-scale information, such as might be used to generate “atlas”-style distribution dot-maps (27), and information on composition and change at the plant community scale, appear to be largely absent at the time of writing, although continuous monitoring of Red Data book plants is in place (Tsintides et al., 2007). Habitat mapping and conservation assessment takes place largely within the Natura 2000 network for areas in the Republic of Cyprus and the Sovereign Base Areas. The frequency of monitoring is strongly associated with reporting obligations and available funds. In addition, biodiversity data are collected as part of research projects, academic projects and environmental impact studies.

For these reasons, experts from across conservation and ecology came together to discuss biodiversity monitoring needs for Cyprus, and to prioritise these according to perceived importance during an expert-elicitation workshop in August 2017. The workshop followed published expert-elicitation methods (44, 56), adhering to the ten guiding principles later published by Roy, Peyton (41), to generate a prioritised list of biodiversity monitoring needs. This is the first time, to our knowledge, that such an approach for prioritising biodiversity monitoring needs has been undertaken within a mediterranean-climate zone.

## Methods

The study was designed to develop a list of prioritised biodiversity monitoring needs for Cyprus. Expert-elicitation methods used by Pocock, Newson (44), to develop a list of attributes for designing biodiversity monitoring programmes in the UK, were adapted for use in Cyprus. The expert-elicitation process for this workshop was carried out using a three-step process (Fig 1):

**Fig 1.**
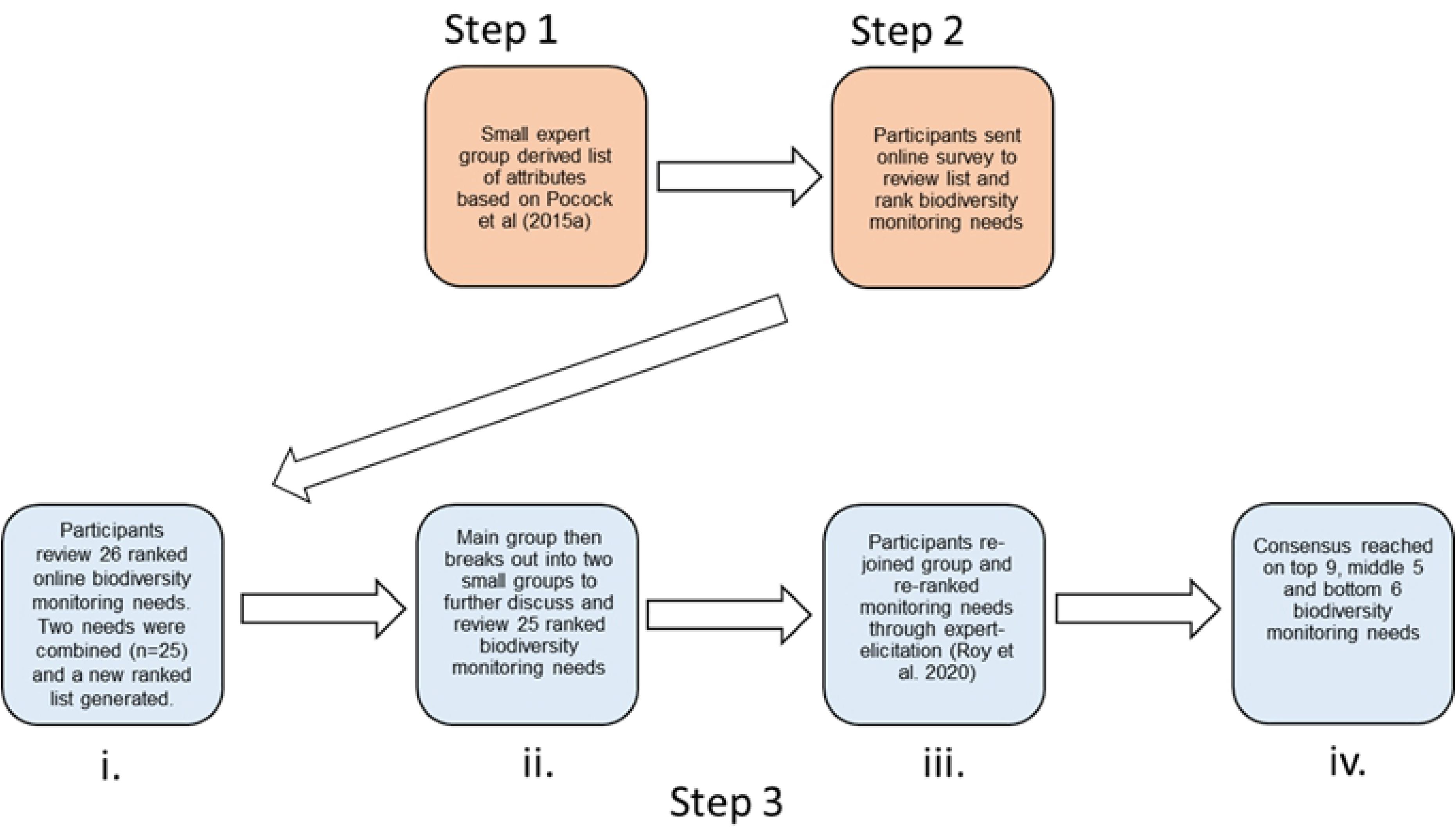
Outline of process to create and prioritise a prioritised consensus list. The process used to create and prioritise a consensus list of biodiversity monitoring needs for Cyprus prior to and during an expert-elicitation workshop in August 2017. Orange boxes denote the work undertaken in advance of the workshop and blue boxes denote tasks undertaken during the workshop. Forty-seven stakeholders were invited to take part in Step 2 of the process; 27 took part. Thirty-nine out of 56 invited stakeholders took part in Step 3 of the process. The 39 stakeholders included those asked to take part in Step 2 but a wider pool of experts were also approached to take part as interest in the workshop increased.

### Step 1: Developing the list of biodiversity monitoring needs in Cyprus

A list of 25 monitoring needs, taken from Pocock, Newson (44), was reviewed by JP (author), Angeliki Martinou and Helen Roy for relevance to Cyprus. The 25 statements were reviewed and the statement text “*There is good retention of contributors”* modified slightly so as to generate a clearer statement for an international audience: *“There is sustained participation”*. The proposed list was then shared with collaborators at the Department of Environment, Government of the Republic of Cyprus, and, after consultation with MH (author), supplemented with an additional monitoring need specific to Natura 2000 sites “*There is extra effort on protected areas, e.g. Natura 2000 sites*”. This resulted in 26 monitoring needs statements for inclusion in the questionnaire described below.

### Step 2: Assessing the importance of the biodiversity monitoring needs for Cyprus

A questionnaire (see SI1), written in both Greek and English, and hosted on the GDPR-compliant survey platform https://www.onlinesurveys.ac.uk, was distributed by email to 47 stakeholders with experience in recording and/or monitoring biodiversity in Cyprus. These stakeholders either had strategic oversight of monitoring, extensive practical experience, and/or participated in monitoring in a professional or voluntary capacity. Stakeholders included volunteer experts who ran biological recording schemes and coordinated others to gather species records, academics from universities and research institutes, and employees from private research and consultancy organisations. They also included stakeholders involved in data collection from government agencies and non-governmental conservation organizations. Those invited possessed expertise in a wide range of taxa and habitats across Cyprus (Table 1, Fig 2).

**Table 1.** Results of ranking exercise of biodiversity monitoring needs statements. Results of ranking exercise of biodiversity monitoring needs statements from an online questionnaire sent to 47 invited stakeholders working in the field of biodiversity monitoring in Cyprus (Step 2 in the expert-elicitation process). Twenty-seven stakeholders ranked their biodiversity monitoring needs for Cyprus from a list of 26 attributes. Stakeholders were asked to rank the statements that represented the 10 most important gaps or opportunities in biological recording in Cyprus, based on their perspective or experience (whereby 1 was the most important gap or opportunity and 10 was the least important gap or opportunity). The number of times each biodiversity monitoring need was selected by all the stakeholders is given alongside the monitoring need statement. This score was then broken down into the number of responses from each of the six stakeholder affiliations and the percent of times it was chosen by the stakeholders within that affiliation, e.g. two consultants and they both selected “There is standardised methodology and protocols to ensure consistency” = 2 (100%). The cells are coloured in a grey-scale continuum, for the percent chosen from within each of the affiliations: 0%, 1-24%, 25-49%, 50-74% and 75-100%. *Themes were added post-workshop.

| * Theme | Step 2 list of biodiversity monitoring need statements | Total<br>number of<br>times<br>selected in<br>top 10 | Biological<br>recorder<br>(n=2) | Consultant<br>(n=2) | Government<br>(n=7) | NGO (n=5) | Researcher<br>(n=9) | Other<br>expertise<br>(n=2) |
| --- | --- | --- | --- | --- | --- | --- | --- | --- |
| Design | There is standardised methodology and protocols to ensure consistency | 18 | - | 2 (100%) | 4 (57%) | 4 (80%) | 7 (78%) | 1 (50%) |
| Human resource | Mentoring, training and support for contributors is provided | 18 | 2 (100%) | 1 (50%) | 5 (71%) | 3 (60%) | 5 (56%) | 2 (100%) |
| Design | There is national or regional co-ordination | 17 | 2 (100%) | 1 (50%) | 5 (71%) | 3 (60%) | 4 (44%) | 2 (100%) |
| Human resource | There are quality assurance checks undertaken in order to ensure the accuracy of the records | 17 | 2 (100%) | 1 (50%) | 4 (57%) | 3 (60%) | 6 (67%) | 1 (50%) |
| Human resource | There are sufficient contributors with specialist knowledge of their taxa | 15 | 1 (50%) | 1 (50%) | 4 (57%) | 3 (60%) | 5 (56%) | 1 (50%) |
| Analytical | There are appropriate analytical/statistical approaches to measure trends from monitoring data | 14 | - | 1 (50%) | 2 (29%) | 4 (80%) | 5 (56%) | 2 (100%) |
| Technological | There are data systems (e.g. online) for efficient data capture and storage | 14 | 1 (50%) | 2 (100%) | 3 (43%) | 2 (40%) | 5 (56%) | 1 (50%) |
| Human resource | There is sustained participation | 12 | 2 (100%) | - | 2 (29%) | 2 (40%) | 5 (56%) | 1 (50%) |
| Design/Human resource | There is wide coverage across the country/region, e.g. covering remote and well-populated areas | 11 | 1 (50%) | - | 1 (14%) | 3 (60%) | 5 (56%) | 1 (50%) |
| Design | There are suitable field sampling methods that are accurate/efficient | 11 | - | 1 (50%) | 4 (57%) | 3 (60%) | 2 (22%) | 1 (50%) |
| Other resource | There are suitable and accessible identification guides | 11 | 2 (100%) | 1 (50%) | 3 (43%) | 1 (20%) | 3 (33%) | 1 (50%) |
| Analytical | Change is reported at appropriate intervals | 10 | - | 1 (50%) | 1 (14%) | 2 (40%) | 6 (67%) | - |
| Human resource | There are sufficient contributors | 10 | 2 (100%) | - | 5 (71%) | 1 (20%) | 1 (11%) | 1 (50%) |
| Design | 'Important' or 'indicator' species have been identified | 10 | - | - | - | 3 (60%) | 5 (56%) | 2 (100%) |
| Analytical | There is access to analytical expertise to measure trends from monitoring data | 10 | - | - | 3 (43%) | 1 (20%) | 5 (56%) | 1 (50%) |
| Human resource | There is appropriate feedback to participants on survey results and findings | 9 | 2 (100%) | 1 (50%) | 2 (29%) | 1 (20%) | 2 (22%) | 1 (50%) |
| Design | Communicate the objectives of monitoring | 9 | 2 (100%) | 2 (100%) | 2 (29%) | 2 (40%) | 1 (11%) | - |
| Design/Human resource | There is extra effort on protected areas, e.g. Natura 2000 sites | 9 | - | - | 2 (29%) | 3 (60%) | 3 (33%) | 1 (50%) |
| Design | There is a scientific scheme design (such as stratified or randomised site selection) for statistical rigour | 8 | - | - | 1 (14%) | 2 (40%) | 3 (33%) | 2 (100%) |
| Analytical | Change is reported on an annual basis | 7 | - | 2 (100%) | 1 (14%) | - | 3 (33%) | 1 (50%) |
| Design/Human resource | There is extra effort on priority species and habitats | 6 | 1 (50%) | 1 (50%) | 1 (14%) | 2 (40%) | 1 (11%) | - |
| Design/Human resource | Recorders collect supplementary data (such as characteristics of the habitat, soil or weather) | 5 | - | - | - | 1 (20%) | 3 (33%) | 1 (50%) |
| Technological/<br>Human resource | The data from monitoring schemes are widely disseminated | 5 | - | 1 (50%) | 1 (14%) | - | 3 (33%) | - |
| Design/Technological | There are simple ways for everyone to report widespread/common/easily-identified species | 4 | - | - | 2 (29%) | 1 (20%) | 1 (11%) | - |
| Human resource | Examples of best practice are identified and shared between schemes and organisations | 4 | 1 (50%) | - | 1 (14%) | 1 (20%) | - | 1 (50%) |
| Technological | There are systems for electronically capturing data in the field | 2 | - | 1 (50%) | 1 (14%) | - | - | - |

**Fig 2.**
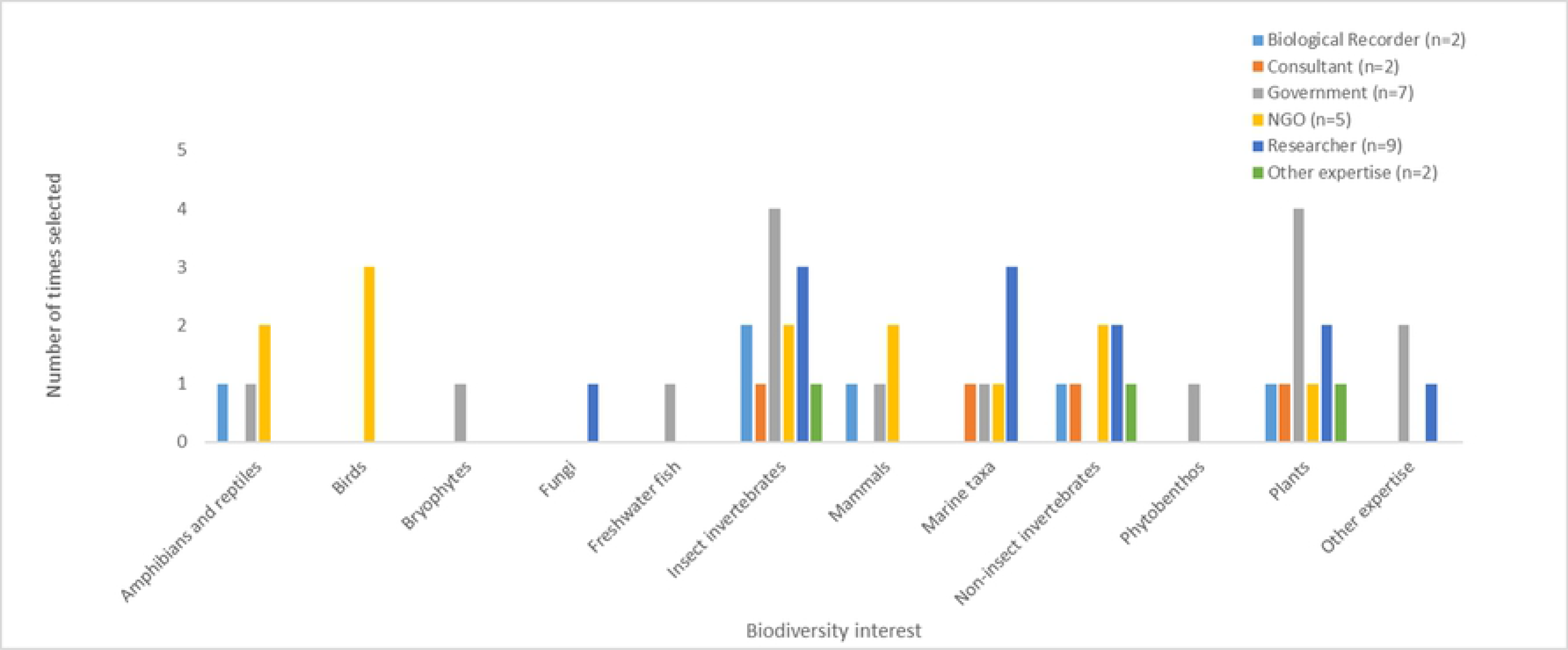
Biodiversity interests of stakeholders. Biodiversity interests of stakeholders working in the field of biodiversity monitoring in Cyprus, who took part in an online questionnaire (Step 2 in the expert-elicitation process); 27 usable responses were received. A pre-populated list of taxonomic interests was given in the question, along with a free text box for additional taxonomic or environment-focused interests. Twenty-three out of the 27 stakeholders selected their top three biodiversity interests; three stakeholders selected more than three biodiversity interests. One stakeholder did not respond and this result was added to “Other expertise”.

The questionnaire was sent to stakeholders two weeks prior to the workshop. Stakeholders were asked to select their organisation from a pre-populated list consisting of the following options: NGO; Government employees; University; Research Institute; Biological Recording (i.e., amateur experts, or citizen scientists); and, Other. Stakeholders were asked to provide information if they selected the “Other” category.

Stakeholders were then asked to select their top three biodiversity interests through the question: “*Please give your top three monitoring / recording priorities. These can either be taxonomy-specific (e.g. monitoring invasive species, birds or butterflies), or environment focused (e.g. monitoring the water levels in the Akrotiri Lake).*” A pre-populated list of taxonomic expertise categories was given within the question (see S1 Table 1), along with a free text box to add other items as needed. Next, stakeholders were asked to select their top ten monitoring needs from the list of 26 provided and to rank them. Stakeholders were encouraged to undertake this ranking task in a two-step process: (1) first review the list of 26 needs and mark ‘N/A’ for the 16 statements that they considered least important, until they were left with the ten that were most important to them; (2) second, assign a score of 1-10 (with 1 being most important and 10 being least) to each of their remaining ten biodiversity monitoring needs. Stakeholders were also invited to add any further comments. These comments were to be picked up in the wider discussion session within the expert-elicitation workshop (Step 3). The results of the ranking exercise from each of the 27 stakeholders who responded (out of the 47 contacted) were combined to generate a single ranked list of 26 biodiversity monitoring needs (Table 1).

The final question asked stakeholders to give one overall score to whether the list of 26 biodiversity monitoring needs adequately represented perceived gaps in Cyprus. Stakeholders were asked to give a score between 1 (not useful at all) to 10 (very useful). SI2 gives the response data received from the 27 stakeholders.

### Step 3: Collaborative prioritisation of an overall list of biodiversity monitoring needs in Cyprus

An expert-elicitation workshop was held on 31st August 2017 for a wider stakeholder network in order to try and reach a consensus on the top biodiversity monitoring needs. Thirty-nine out of 56 invited stakeholders attended the workshop. These stakeholders were from the same sectors as in Step 2, and were from both the UK and Cyprus; UK participants were either active in biodiversity recording in Cyprus, or were invited to provide their experiences around the reality of prioritising particular biodiversity monitoring infrastructures with limited resources. The 39 stakeholders included some of those who had taken part in Step 2 (n = 16) and also stakeholders who had not (n = 23). The workshop was combined with a plenary session with presentations from all stakeholder groups, in order for the stakeholders to understand the range of monitoring already undertaken across Cyprus, and to learn about relevant UK schemes and infrastructure. An overview presentation of the summary results from the online survey was also given during the plenary session (SI2).

#### Step 3i

Following the presentation of the online survey results, stakeholders discussed and reviewed the ranked list of 26 biodiversity monitoring needs from the online survey (Step 2) during the plenary. During the discussions, it was agreed that two of the monitoring needs could be merged: “*There are sufficient contributors with specialist knowledge of their taxa combined”* was merged with “*There are sufficient contributors”*, to give: *There are sufficient ‘and sustained’ contributors with specialist knowledge of their taxa”*. This then gave 25 biodiversity monitoring needs statements taken forward for further discussion and ranking in breakout groups.

#### Step 3ii

The group of 39 stakeholders was divided into two breakout groups to discuss and review the ranked list of the 25 biodiversity monitoring needs. The breakout groups then re-merged to review the biodiversity monitoring needs and generate an agreed overall list of these.

#### Step 3iii

The results (S2) of the online ranking exercise of biodiversity monitoring needs were summarised (Table 1) and used as a point from which to start the consensus-building process. During the exercise it was made clear to participants that each biodiversity monitoring need could be moved up or down the list, with the ultimate location in the list not being directly dependent on the original online rank. Stakeholders were also encouraged to edit the needs statements where it was felt they needed combining, or exhibited redundancy, and also to suggest additional needs that were missing from the original list.

#### Step 3iv

The needs statements were collectively prioritised into three categories during the expert-elicitation process, the top (most important), middle (intermediate importance) and bottom (least important) statements (Table 3).

After the workshop, general themes were ascribed to all the needs statements in order to assess the relative priorities of different types; however, these should not be taken as the only possible categorisation, and hybrid themes were also used to recognise that some needs combined elements of more than one theme (Tables 1 and 3, Fig 3). Five themes were assigned as follows: Analytical, e.g. the capacity to analyse data to produce biodiversity time trends; Design, e.g. monitoring design for the programme including protocols around reporting of species; Human resource, e.g. the time contribution or availability of the participant or coordinator to ensure that data are collected and collated; Technological, e.g. the use of methods involved in data capture such as data loggers or the use of mobile applications; and Other resource, e.g. any resource that was not considered to fit within the previous themes. A sixth theme was assigned to the additional comments received as part of the online questionnaire and expert-elicitation workshop, “Political”. This theme was assigned to comments where overarching or management-level decision making issues were raised. Additional comments raised from the online survey and workshop were brought into the discussions of the expert-elicitation workshop, but these points were not ultimately included in the final list.

**Fig 3.**
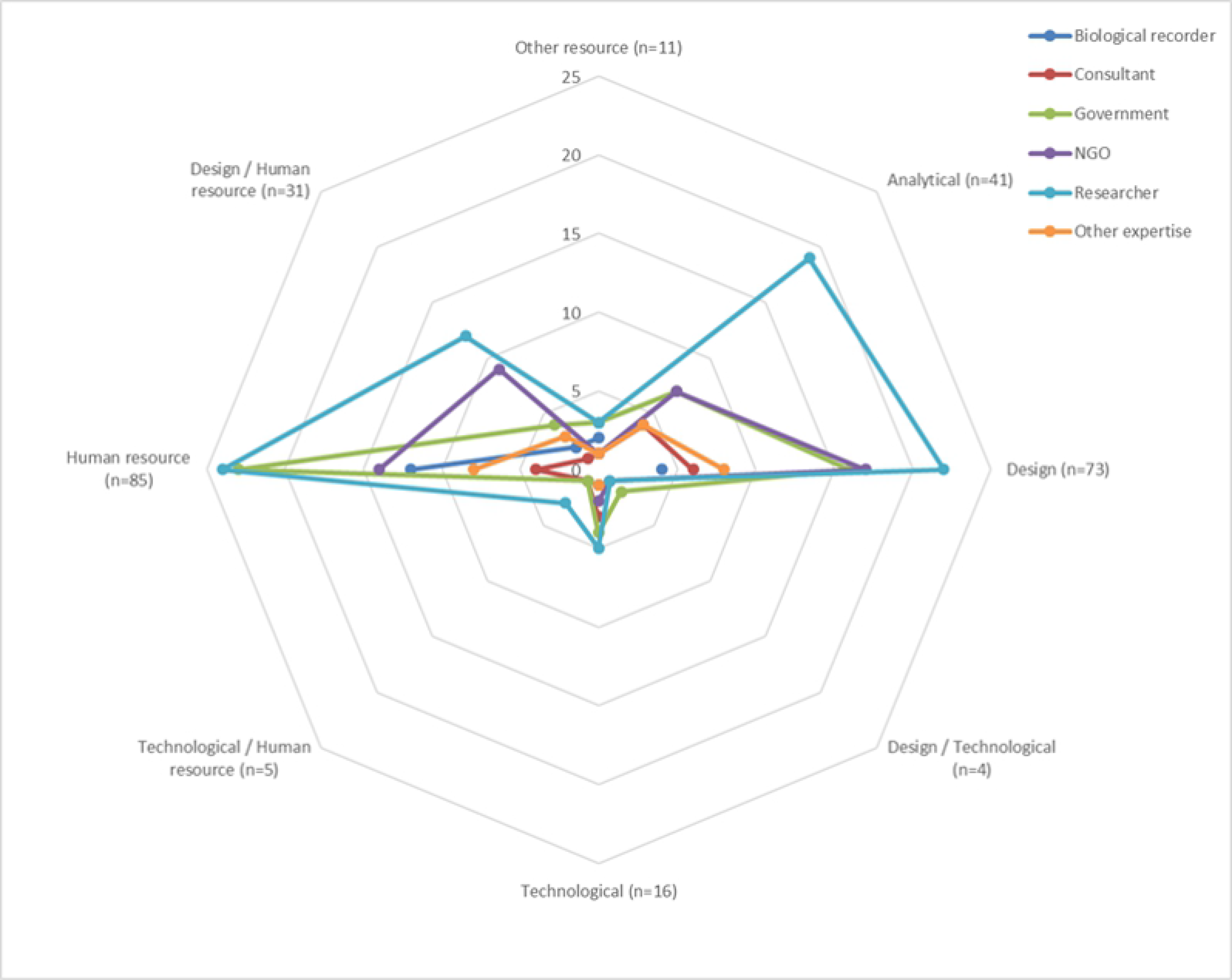
Radar chart showing the results of an online questionnaire. Radar chart showing the results of an online questionnaire sent to 47 stakeholders working in the field of biodiversity monitoring in Cyprus. The 26 biodiversity monitoring need statements were resolved into eight themes post-workshop: Analytical, Design, Design/Technological, Technological, Technological/Human resource, Human resource, Design/Human Resource, and Other resource. The number of times stakeholders selected each of the eight themes is given on the radial axis next to the theme name.

The individual themes were combined where biodiversity monitoring needs were considered to have spanned several themes, e.g. Design/Human resource for the need: “*There is wide coverage across the country/region, e.g. covering remote and well-populated areas*” as to realise this need would require both participant engagement to physically record in different areas, alongside survey design to ensure representative distribution data were collected.

## Results

### Step 2. Online survey results

#### Stakeholder affiliation

Twenty-nine out of 47 invited experts completed the online questionnaire prior to the workshop; two results were excluded due to not having sufficient data for analysis, leaving 27 records for use in the workshop (SI2). The 27 stakeholder respondent affiliations were re-categorised post-workshop, based on the information provided, in order to create what were considered to be more informative groups. As such, University and non-private Research Institute were combined to give “Researcher”, and “Consultant” was added to give more detail for affiliations previously listed as “Other”. The following six post-workshop affiliation types were used in the analyses: Biological recorders (n = 2); Consultants (n = 2); Government staff (n = 7); NGO staff (n = 5); Researcher (n = 9); and Other expertise (including non-response) (n = 2).

#### Stakeholder biodiversity interests

Fig 2 provides a summary of the results from stakeholders who responded to the online question asking for information on biodiversity interests. Note that the category “Insect invertebrates” included those with expertise in aquatic invertebrates; also, “inland fish” was renamed “freshwater fish” for clarity. “Other expertise” included non-response data. No environment-focused responses were given (Fig 2). A maximum of 81 affiliation/biodiversity interest combinations were possible for the 27 respondents (three choices x 27). Fifty-four combinations were returned, with three respondents choosing more than three options. Government and research stakeholders showed a broad range of monitoring interests across taxonomic groups, while the NGO stakeholders focused on amphibians, reptiles and birds. Plants and insect invertebrates (including aquatic taxa) were the best represented taxonomic groups (Fig 2).

#### Top ten priority monitoring needs

The results of the online ranking exercise (Step 2) for the top ten needs are given in Table 1. All 26 monitoring needs from the original list were selected at least once by a stakeholder. The attributes “*There is standardised methodology and protocols to ensure consistency*” and *“Mentoring, training and support for contributors is provided “* were the two top selections (each was selected 18 times) from the online surveys. Monitoring needs primarily relating to reporting (i.e. recording), such as “*There are simple ways for everyone to report widespread / common / easily-identified species*” and “*There are systems for electronically capturing data in the field*”, along with “*Examples of best practice are identified and shared between schemes and organisations*” were ranked lowest by the respondents (Table 1). A maximum of 270 top ten selected statements was possible (27 respondents x 10 statements), and a total of 266 selections were actually made. Additional comments on the survey questions, or points considered important, captured from stakeholders are given in SI3.

There was broad agreement between and within affiliation type for the higher ranking statements (Table 1), with agreement generally decreasing further down the list of biodiversity monitoring needs. The biodiversity need: “*There is standardised methodology and protocols to ensure consistency*” was considered to be of importance by Consultants, Government representatives, NGOs and Researchers, with over half the respondents within these affiliations selecting it as one of the top ten needs. Biological recorders (i.e. amateur experts) selected a narrower range of monitoring needs that they considered to be of importance, with 13 out of a possible twenty statements selected; this is in comparison to Consultants and Other expertise who selected 17 and 18 out of 26 respectively.

The two most important themes, with importance being assigned by the number of times they were placed in the top 10, were Design and Human resource (Fig 3). Design, Human resource and Design/Human Resource-based needs statements were selected 189 times out of the 266 selections made. Needs classified as Analytical were selected the second most frequently (n = 41). Technological and other (non-human) resource-based needs were viewed as less important by stakeholders.

Overall scoring for the adequacy of the survey needs statements in representing biodiversity monitoring gaps in Cyprus was favourable (Table 2). The average score per affiliation-type was over seven (with 10 being most useful). “Governmental” stakeholders agreed most strongly with the statements (average score 9.2), and the variability in the range of scores was generally low across the stakeholder groups, with the exception of the “Researcher” affiliation that had scores ranging from 1 through to 10. The lowest average score was for the “Researcher” category at 7.4 out of 10.

**Table 2.**
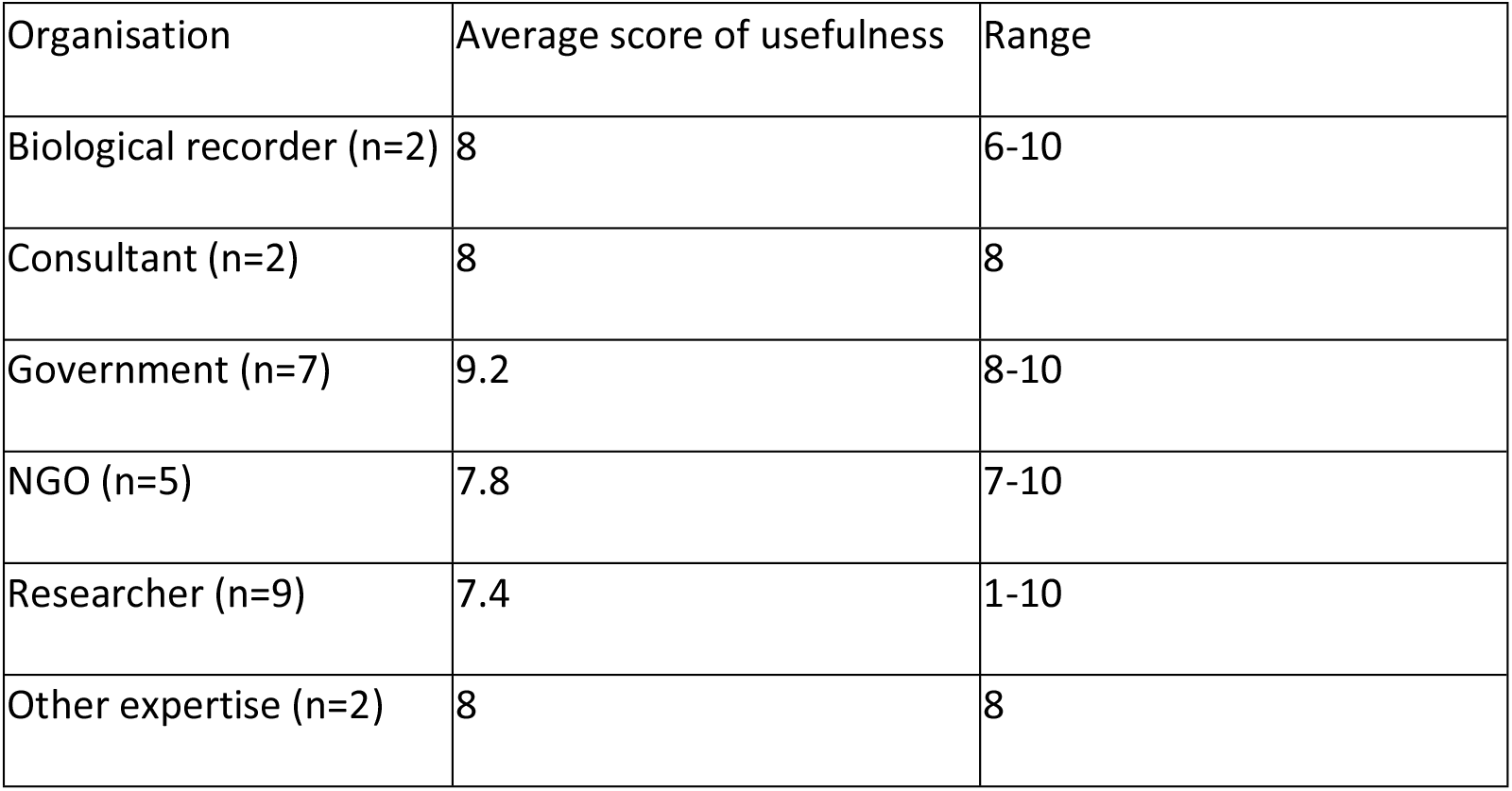
Results of stakeholder responses. Results of 26 stakeholders responses asked whether the online questionnaire (S1 Table 1) adequately represented any gaps in biological monitoring in Cyprus. Stakeholders were asked to score how useful the survey was from 10 ‘very useful’ to 1 ‘not useful at all’.

### Step 3. Expert-elicitation workshop

3.i. Twenty-five monitoring needs were taken forward for voting by stakeholders whilst together in plenary, following the merging of two statements, as outlined in the methods above.

3.ii and 3.iii. During the expert-elicitation process that followed the breakout group discussions, the 25 needs were further modified and combined in order to better align with the stakeholders needs or to reduce redundancy within the statements. The following needs were modified:

1. *“There is standardised methodology and protocols to ensure consistency”*. This was amended to incorporate the following four needs:

- *There are suitable field sampling methods that are accurate/efficient*
- *There are appropriate analytical/statistical approaches to measure trends from monitoring data*
- *Recorders collect supplementary data (such as characteristics of the habitat, soil or weather).* This need was considered important for the marine environment as certain data, e.g. temperature or depth, were considered hard to get for this environment
- *Examples of best practice are identified and shared between schemes and organisations*
2. *“Change is reported at appropriate intervals”* and *“Change is reported on an annual basis”* were combined to give the new biodiversity monitoring need “*Change is reported at appropriate intervals e.g. on an annual basis”*.
3. *“There are data systems (e.g. online) for efficient data capture and storage”* biodiversity monitoring need was amended to *“There are improved and accessible data systems (e.g. online) for efficient data capture and storage”*.

During the expert-elicitation workshop, separate needs statements were not assigned individual ranks, but rather consensus was reached on the relative positioning of groups of statements taken together. The following ranked groups were established: a top priority group of nine statements; a middle group of five statements; and a bottom group of six statements (Table 3). The ranked positions of the needs statements from the online survey were very similar to those decided upon during the workshop (accounting for the editing of the six statements described previously), with statements broadly being found in the same relative positions in terms of top, middle and lower levels (Table 3). Eight of the top nine biodiversity monitoring needs from the workshop were also in agreement with the results of the online survey (Tables 1, 3). The largest change in priority for a need between the online survey and the workshop was *“Communicate the objectives of monitoring”* which increased in relative importance during the workshop, having been selected only nine times during the online scoring exercise but becoming a top need in the workshop.

**Table 3.** Consensus-ranked list of biodiversity monitoring needs. Consensus-ranked list of biodiversity monitoring needs from the expert-elicitation workshop (Step 3), held in Cyprus in August 2017. Thirty-nine stakeholders participated in the workshop from the field of biodiversity monitoring in Cyprus and the UK. Stakeholders considered the results from the online survey, in two smaller breakout groups. The number of votes each monitoring need received in the online survey was given and was used as a method for initially ranking the needs at the start of the elicitation work. During the expert-elicitation process, five (of the 25) monitoring needs were combined with other statements to generate a final list of 20 biodiversity monitoring needs for Cyprus. Consensus was reached on the top nine statements, the middle five statements and the bottom six statements. Themes were added post-workshop.

| Theme | Step 3iii List of biodiversity monitoring need statements | Consensus priority group |
| --- | --- | --- |
| Human resource | Mentoring, training and support for contributors is provided | Top |
| Design | There is standardised methodology and protocols to ensure consistency | Top |
| Human resource | There are quality assurance checks undertaken in order to ensure the accuracy of the records | Top |
| Design | There is national or regional coordination | Top |
| Human resource | There are sufficient 'and sustained' contributors with specialist knowledge of their taxa | Top |
| Technological | There are improved and accessible data systems (e.g. online) for efficient data capture and storage | Top |
| Human resource | There is sustained participation | Top |
| Design/Human resource | There is wide coverage across the country/region, e.g. covering remote and well-populated areas | Top |
| Design | Communicate the objectives of monitoring | Top |
| Other resource | There are suitable and accessible identification guides | Middle |
| Analytical | Change is reported at appropriate intervals e.g. on an annual basis | Middle |
| Design | 'Important' or 'indicator' species have been identified | Middle |
| Human resource | There is appropriate feedback to participants on survey results and findings | Middle |
| Analytical | There is access to analytical expertise to measure trends from monitoring data | Middle |
| Design/Human resource | There is extra effort on protected areas, e.g. Natura 2000 sites | Bottom |
| Design | There is a scientific scheme design (such as stratified or randomised site selection) for statistical rigour | Bottom |
| Design/Human resource | There is extra effort on priority species and habitats | Bottom |
| Technological/Human resource | The data from monitoring schemes are widely disseminated | Bottom |
| Design/Technological | There are simple ways for everyone to report widespread/common/easily-identified species<br>[wider engagement] | Bottom |
| Technological | There are systems for electronically capturing data in the field | Bottom |

## Discussion

Globally there is a need to deliver robust data to quantify changes in biodiversity, thereby helping to ensure that conservation funds and other resources are used optimally. This goal, however, also requires an appropriate prioritisation strategy wherever resources are limited, e.g. in a given region should we focus on training volunteers or employing more professionals? Should we record broad-scale distributions across multiple taxa, or intensively monitor a small selection of species’ populations? To assist with this goal, we have developed an ordered set of biodiversity monitoring needs for Cyprus using questionnaire-based online rankings and subsequent workshop-based expert-elicitation. High priority needs in Cyprus primarily focused around two themes: Human resource and Design. Despite technologies playing an important role in advancing the generation and utilisation of biological data for conservation in some locations (August et al. 2015), we found that technology-based needs were a lower priority for our stakeholders. Needs related to data analysis ranked in the middle of both our online and expert-elicitation rankings.

Monitoring needs focused on design were prioritised highly, but those needs focused on human resources, such as “*Mentoring, training and support for contributors is provided*”, “*There are quality assurance checks undertaken in order to ensure the accuracy of the records*” and “*There are sufficient ‘and sustained’ contributors with specialist knowledge of their taxa*”, were also important. Volunteer enthusiasm is known to be a key driver in sustaining participation in biodiversity recording schemes (44), and this can be supported through mentoring and training (57).

Expert-elicitation, as a mechanism for the development of a prioritised list of biodiversity monitoring needs in Cyprus, was selected following the successful application of this technique in the UK (44). Despite the many recorded benefits of expert-elicitation, this approach to elucidating answers can risk bringing in participant bias (40). Moreira, Allsopp (58) recommend that alongside increasing transdisciplinary research, the identification of research priorities needs to be inclusive of the range of stakeholder interests, as well as regional needs and possible funding mechanisms. Within our workshop, although 27 stakeholders took part in the online survey and 39 stakeholders took part in the workshop, the biological recording community (i.e. amateur experts, or citizen scientists collecting biological records) was arguably under-represented, which may have shifted the focus of the prioritisation results towards data user needs, as data contributors can have different motivations to data users. That the stakeholders participating in both the online questionnaire and expert-elicitation workshop were primarily end-users of data, and yet did not solely prioritise needs focused on design and analysis (professional end users typically rely on detailed and accurate systematic data to inform decision making), could be attributed to limitations inherent to one often encountered definition of data users. Data users are traditionally considered to be academics or government agencies (44). Whilst this assumption may hold true in some places and at some times, it may be the case that data users are also contributors of biodiversity data. Those stakeholders who are also undertaking recording in a voluntary capacity may therefore have a more human resource-focused view of monitoring needs. As such, through this expert-elicitation process, we suggest that we have reached a reasonably robust community consensus on the biodiversity monitoring needs most relevant to Cyprus.

It was unexpected that technology-based needs, and needs associated with technology, were not ranked consistently highly in both the online questionnaire and workshop. Advances in technology, for example through websites that allow for rapid data entry, promote straightforward data sharing, or which facilitate the automation of record validation, are an important part of modern biological recording (59). However, the ready availability of some technologies at this point in history, such as mobile applications and data sharing portals, are likely to mean that these biodiversity monitoring needs are not as important when compared to having enough human resources (support and mentoring etc.) to undertake the recording in the first instance.

Reporting back to data collectors is often seen as a key motivator for ensuring engagement of recorders (e.g. 24). That the biodiversity monitoring need “*The data from monitoring schemes are widely disseminated*” was one of the lowest priorities for both the online questionnaire and expert-elicitation workshop was also unexpected. Its placement could be due to other biodiversity monitoring needs being considered higher priorities, rather than this aspiration being unimportant *per se*. It could also be due to local scientific monitoring cultures in place on the island that resist data sharing due to perceptions relating to the loss of control over such data, and/or the lack of professional reward currently associated with such actions, as was suggested by one participant in our workshop.

The successful adaptation of the framework of Pocock, Newson (44) supports their contention that the framework was replicable and robust for other countries. With respect to the prioritisation of the biodiversity monitoring needs, our results broadly reflected the findings of Pocock, Newson (44), with both the online and expert-elicitation workshop prioritising standard methodologies, quality assurance, and suitable field sampling methods that are accurate and efficient. Our findings did however deviate in the increased prioritisation of needs relating to national/regional coordination, and the collection of additional environmental data, such as climatic conditions and additional data for the marine environment, which were listed in the top nine biodiversity monitoring needs in the current study. They were amongst the least important in Pocock, Newson (44). The collection of additional environmental data, such as habitat information, can increase the value of records in answering ecological questions (60). The relative importance here may in part be due to concern over the impacts of climate change on biodiversity, as the Mediterranean is predicted to be affected directly through climatic conditions such as drought and high temperatures (61, 62). The specific reference to collecting data in the marine environment is likely linked to both the lack of data in many taxonomic groups of marine organisms and the lack of available environmental data for this in Cyprus. The Mediterranean Sea is facing multiple pressures from climate change, overexploitation of marine resources and IAS (63), and, as such, additional data to monitor and record change would be important in this vulnerable system.

### Possible uses and next steps for the prioritised list of biodiversity monitoring needs

#### Informing the design of monitoring schemes

Pocock, Newson (44) recommend using such prioritised lists to look for gaps in existing programmes, and also in the design of new biodiversity monitoring programmes. We found that the most highly prioritised biodiversity monitoring needs were those related to the development of robust methodologies and ensuring a suitable geographic spread of sufficiently skilled and informed contributors. These prioritised needs could be used to help focus existing resources for data collection. For example, supporting the development of standardised methodologies, thus ensuring consistency in the way records are collected in new schemes, and recognition of the importance of mentoring, training and supporting volunteers in existing, or newly establishing, schemes which could result in increased funding for these activities (57).

#### Increasing infrastructure capacity

National coordination was also a top biodiversity monitoring need (Table 3). Although there are many biodiversity datasets within Cyprus, there are gaps in collating, analysing, identifying trends and needs, and communicating and sharing data. There would be merit in developing guidance on mechanisms for extracting data from a selection of sources (e.g. from studies undertaken for another purpose, but that include biodiversity data), how to manage this information (e.g. existence of appropriate software and training to use it), and how to analyse, interpret, and share it among the interested parties (13).

Establishing a national hub for the centralised collation, dissemination and analysis of biodiversity monitoring data could help to deliver the data needed to enable governments to report on environmental change, and help to prioritise suitable monitoring approaches for sites or species that need protecting, or for which monitoring is specifically mandated by law (64). A national hub holding high quality, verified biological records could also play a role in coordinating and supporting a network of citizen science recorders, a model successfully employed in the UK for over 50 years (64). Such a national hub was suggested from the online survey by one respondent (SI3) and could be a valuable tool in delivering knowledge about biodiversity in Cyprus. Developing a GBIF Participant node could help enable the coordination of networks of people and institutions that create, manage and utilise data on biodiversity across Cyprus. However, there can be mistrust from data users over the quality of the data in large aggregated datasets, such as inconsistent taxonomy, inadequate spatial coverage, or lack of reported protocols (65), which can lead to reduction in the use of the data for scientific studies (66). Franz and Sterner (2018) also argue for greater accountability of the role of these databases in making choices that directly underpin the perceived quality of aggregated biodiversity data, such as the creation and maintenance of taxonomic hierarchies that may not reflect the concepts used in local biodiversity communities; these authors call for global data hubs to work to develop new technical pathways and social incentives as mechanisms to bridge the current trust gap.

#### Developing cross-taxa networks

“*Examples of best practice are identified and shared between schemes and organisations*” was not highly scored during the online questionnaire (Table 1). During the expert-elicitation workshop, this biodiversity monitoring need was subsequently amalgamated with three other needs: “*There are suitable field sampling methods that are accurate/efficient*”, “*There are appropriate analytical/statistical approaches to measure trends from monitoring data*” and “*Recorders collect supplementary data (such as characteristics of the habitat, soil or weather)*” within the need “*There is standardised methodology and protocols to ensure consistency”*. This was one of the top nine biodiversity monitoring needs. Should this need become a higher priority on Cyprus, the approach taken by the National Forum for Biological Recording (NFBR) charity in the UK may be relevant. The NFBR works across taxonomic groups to support biological recorders, hosts conferences, and represents their interests in governmental decision-making, through mechanisms such as the State of Nature report (67). Such a model could enable the Cypriot biological recording community to share ideas and experiences for the maintenance and support of recording.

## Conclusion

Prioritising biodiversity monitoring needs is becoming increasingly important as we continue to see major declines in biodiversity and degradation of ecosystem function (1). Twenty biodiversity monitoring needs were successfully prioritised for Cyprus during an expert-elicitation workshop, with equal priority assigned to different monitoring needs within three categories: a top nine, a middle five, and a bottom six. We hope that this prioritised list of needs will assist with the delivery of robust monitoring programmes that result in both high-value biodiversity monitoring data for end users and support the motivations and values of the participants.

## Acknowledgements

The authors would like to thank all the participants who took part in the online questionnaire and workshop. The authors would also like to thank the Akrotiri Environmental Education Centre for hosting the workshop and for their continued support for our project, Dr Angeliki Martinou for her initiation of the work and assistance with aspects of the methodology, and Professor Helen Roy for her role in facilitating the expert-elicitation process.

## Supporting information

S1 Table 1. **Online survey questionnaire.** English version of online survey questionnaire sent to stakeholders working in the field of biodiversity monitoring in Cyprus (in both Greek and English). Results were collected using the online survey platform https://www.onlinesurveys.ac.uk/.

S2 Table 2. **Unprocessed results from an online survey.** Unprocessed results from an online survey sent to stakeholders working in the field of biodiversity monitoring in Cyprus (in both Greek and English). Information that would make individuals identifiable has been removed for the purposes of publication. Each respondent has a unique code generated by the online survey. Respondents were asked if they were able to attend the workshop. They were also asked to select their organisation from a pre-populated list or to select “other” and then specify. Responses that were submitted in Greek were translated by the project partners. Respondents were then asked to give their top three biodiversity monitoring interests from a pre-populated list with a free text box for additional information. Respondents were then asked to rank statements given in a pre-populated list that represent the 10 most important gaps or opportunities in biological recording. Next, stakeholders were asked to rank how adequately the questionnaire addressed the biological recording gaps in Cyprus. Finally, stakeholders were asked if they wished to be co-authors on the current manuscript.

S3 Table 3. **Additional stakeholder comments from both an online questionnaire and workshop.** Additional stakeholder comments from both an online questionnaire and workshop involving stakeholders from the field of biodiversity monitoring in Cyprus. Stakeholders were asked to give further features or comments that they considered important and that might have missed out from either the online survey or that came up as part of the discussions. A facilitator summarised the workshop discussion points. This task was undertaken as part of a wider exercise on ranking the biodiversity monitoring needs for Cyprus. *Themes were added post-workshop.

## References

1. Díaz S, Settele J, Brondízio E, Hien N, M G, Agard J, et al. Summary for policymakers of the global assessment report on biodiversity and ecosystem services of the Intergovernmental Science-Policy Platform on Biodiversity and Ecosystem Services. IPBES, 2019 6 May 2019. Report No.

2. Erotokritou E, Vogiatzakis IN. Landscape linkages for the distribution of the endangered Hierophis cypriensis in Cyprus. Ecologia mediterranea: Revue internationale d’écologie méditerranéenne= International Journal of Mediterranean Ecology. 2019;45(1):31–44.

3. Sala OE, Stuart Chapin F, III, Armesto JJ, Berlow E, Bloomfield J, et al. Global Biodiversity Scenarios for the Year 2100. Science. 2000;287(5459):1770–4. doi: 10.1126/science.287.5459.1770.

4. Coll M, Piroddi C, Steenbeek J, Kaschner K, Lasram FBR, Aguzzi J, et al. The biodiversity of the Mediterranean Sea: estimates, patterns, and threats. PloS one. 2010;5(8):e11842.

5. Hall CM. Tourism and biodiversity: more significant than climate change? Journal of Heritage Tourism. 2010;5(4):253–66. doi: 10.1080/1743873X.2010.517843.

6. Sánchez-Bayo F, Wyckhuys KAG. Worldwide decline of the entomofauna: A review of its drivers. Biol Conserv. 2019;232:8–27. doi: https://doi.org/10.1016/j.biocon.2019.01.020.

7. Nieto A, Roberts SP, Kemp J, Rasmont P, Kuhlmann M, Criado MG, et al. European red list of bees. 2017.

8. Tsikliras AC, Dinouli A, Tsiros V-Z, Tsalkou E. The Mediterranean and Black Sea Fisheries at Risk from Overexploitation. PLOS ONE. 2015;10(3):e0121188. doi: 10.1371/journal.pone.0121188.

9. McNeely JA. The sinking ark: pollution and the worldwide loss of biodiversity. Biodiversity & Conservation. 1992;1(1):2–18.

10. Defra. A Green Future: Our 25 Year Plan to Improve the Environment. Defra, 2018.

11. Gitzen RA, Millspaugh JJ, Cooper AB, Licht DS. Design and analysis of long-term ecological monitoring studies: Cambridge University Press; 2012.

12. Hochkirch A, Samways MJ, Gerlach J, Böhm M, Williams P, Cardoso P, et al. A strategy for the next decade to address data deficiency in neglected biodiversity. Conserv Biol. 2020.

13. Groom QJ, Desmet P, Vanderhoeven S, Adriaens T. The importance of open data for invasive alien species research, policy and management. Management of Biological Invasions. 2015;6(2):119.

14. Beck J, Ballesteros-Mejia L, Nagel P, Kitching IJ. Online solutions and the ‘Wallacean shortfall’: what does GBIF contribute to our knowledge of species’ ranges? Divers Distrib. 2013;19(8):1043–50. doi: 10.1111/ddi.12083.

15. Brown J, Lomolino M. Biogeography, 2nd edn Sinauer Associates: Sunderland. MA, USA. 1998.

16. Brito D. Overcoming the Linnean shortfall: Data deficiency and biological survey priorities. Basic and Applied Ecology. 2010;11(8):709–13. doi: https://doi.org/10.1016/j.baae.2010.09.007.

17. de Carvalho MR, Bockmann FA, Amorim DS, Brandão CRF, de Vivo M, de Figueiredo JL, et al. Taxonomic Impediment or Impediment to Taxonomy? A Commentary on Systematics and the Cybertaxonomic-Automation Paradigm. Evolutionary Biology. 2007;34(3):140–3. doi: 10.1007/s11692-007-9011-6.

18. Kim KC, Byrne LB. Biodiversity loss and the taxonomic bottleneck: emerging biodiversity science. Ecological Research. 2006;21(6):794.

19. Boxshall G, Self D. UK Taxonomy & Systematics Review–2010. Results of survey undertaken by the Review Team at the Natural History Museum serving as contractors to the Natural Environment Research Council (NERC). 2011.

20. Juanes F. Visual and acoustic sensors for early detection of biological invasions: Current uses and future potential. Journal for Nature Conservation. 2018;42:7–11. doi: https://doi.org/10.1016/j.jnc.2018.01.003.

21. Piaggio AJ, Engeman RM, Hopken MW, Humphrey JS, Keacher KL, Bruce WE, et al. Detecting an elusive invasive species: a diagnostic PCR to detect B urmese python in F lorida waters and an assessment of persistence of environmental DNA. Molecular ecology resources. 2014;14(2):374–80.

22. Agnarsson I, Kuntner M. Taxonomy in a changing world: seeking solutions for a science in crisis. Systematic Biology. 2007;56(3):531–9.

23. Hochkirch A. The insect crisis we can’t ignore. Nature. 2016;539(7628):141-.

24. Morris R. A change in funding directions: Implications for biological recording. British Journal of Entomology and Natural History. 2012;25(3):143.

25. Isaac NJB, Pocock MJO. Bias and information in biological records. Biological Journal of the Linnean Society. 2015;115(3):522–31. doi: 10.1111/bij.12532.

26. Kampen H, Medlock JM, Vaux AGC, Koenraadt CJM, van Vliet AJH, Bartumeus F, et al. Approaches to passive mosquito surveillance in the EU. Parasites & Vectors. 2015;8(1):9. doi: 10.1186/s13071-014-0604-5.

27. Preston CD. Following the BSBI’s lead: the influence of the Atlas of the British flora, 1962–2012. New Journal of Botany. 2013;3(1):2–14. doi: 10.1179/2042349713Y.0000000020.

28. Pescott OL, Walker KJ, Pocock MJ, Jitlal M, Outhwaite CL, Cheffings CM, et al. Ecological monitoring with citizen science: the design and implementation of schemes for recording plants in Britain and Ireland. Biological Journal of the Linnean Society. 2015;115(3):505–21.

29. Pocock MJO, Roy HE, Preston CD, Roy DB. The Biological Records Centre: a pioneer of citizen science. Biological Journal of the Linnean Society. 2015;115(3):475–93. doi: 10.1111/bij.12548.

30. Brereton T, Botham M, Middlebrook I, Randle Z, Noble D, Roy D. United Kingdom Butterfly Monitoring Scheme Annual Report 2016. 2017.

31. Comont R, Miles S. BeeWalk Annual Report 2019. Bumblebee Conservation Trust, Stirling, Scotland UK. 2019.

32. CIESM ICfSEotMS. CIESM JellyWatch Program 2014 [cited 2021 12/04/2021]. Available from: http://www.ciesm.org/marine/programs/jellywatch.htm.

33. Crall AW, Renz M, Panke BJ, Newman GJ, Chapin C, Graham J, et al. Developing cost-effective early detection networks for regional invasions. Biol Invasions. 2012;14(12):2461–9. doi: 10.1007/s10530-012-0256-3.

34. Giovos I, Kleitou P, Poursanidis D, Batjakas I, Bernardi G, Crocetta F, et al. Citizen-science for monitoring marine invasions and stimulating public engagement: a case project from the eastern Mediterranean. Biol Invasions. 2019;21(12):3707–21.

35. Groom Q, Strubbe D, Adriaens T, Davis AJ, Desmet P, Oldoni D, et al. Empowering Citizens to Inform Decision-Making as a Way Forward to Support Invasive Alien Species Policy. Citizen Science: Theory and Practice. 2019;4(1).

36. Kleitou P, Giovos I, Wolf W, Crocetta F. On the importance of citizen-science: the first record of Goniobranchus obsoletus (Rüppell and Leuckart, 1830) from Cyprus (Mollusca: Gastropoda: Nudibranchia). BioInvasions Records. 2019;8(2):252–7.

37. Wetzel FT, Bingham HC, Groom Q, Haase P, Kõljalg U, Kuhlmann M, et al. Unlocking biodiversity data: Prioritization and filling the gaps in biodiversity observation data in Europe. Biol Conserv. 2018;221:78–85.

38. García-Barón I, Giakoumi S, Santos MB, Granado I, Louzao M. The value of time-series data for conservation planning. J Appl Ecol. 2021.

39. Colson AR, Cooke RM. Expert Elicitation: Using the Classical Model to Validate Experts’ Judgments. Review of Environmental Economics and Policy. 2018;12(1):113–32. doi: 10.1093/reep/rex022.

40. Sutherland WJ, Burgman M. Policy advice: use experts wisely. Nature News. 2015;526(7573):317.

41. Roy HE, Peyton JM, Booy O. Guiding principles for utilizing social influence within expert-elicitation to inform conservation decision-making. Glob Change Biol. 2020;n/a(n/a). doi: 10.1111/gcb.15062.

42. Sutherland WJ, Dias MP, Dicks LV, Doran H, Entwistle AC, Fleishman E, et al. A horizon scan of emerging global biological conservation issues for 2020. Trends Ecol Evol. 2020;35(1):81–90.

43. Grace M, Balzan M, Collier M, Geneletti D, Tomaskinova J, Abela R, et al. Priority knowledge needs for implementing nature-based solutions in the Mediterranean islands. Environmental Science & Policy. 2021;116:56–68. doi: https://doi.org/10.1016/j.envsci.2020.10.003.

44. Pocock MJ, Newson SE, Henderson IG, Peyton J, Sutherland WJ, Noble DG, et al. Developing and enhancing biodiversity monitoring programmes: a collaborative assessment of priorities. J Appl Ecol. 2015;52(3):686–95.

45. CBD CoBD. Fourth National Report to the United Nations Convention on Biological Diversity. Nicosia: Department of Environment - Ministry of Agriculture, Natural Resources and Environment, 2010.

46. Delipetrou P, Makhzoumi J, Dimopoulos P, Georghiou K. Cyprus. Mediterranean Island Landscapes: Springer; 2008. p. 170–203.

47. Vogiatzakis IN, Manolaki P, Zomeni M, Zotos S. Habitats. In: Sparrow DJ, John E, editors. An introduction to the wildlife of Cyprus: Terra Cypria; 2016.

48. Manolaki P, Zotos S, Vogiatzakis IN. An integrated ecological and cultural framework for landscape sensitivity assessment in Cyprus. Land Use Policy. 2020;92:104336. doi: https://doi.org/10.1016/j.landusepol.2019.104336.

49. Varnava AI, Roberts SP, Michez D, Ascher JS, Petanidou T, Dimitriou S, et al. The wild bees (Hymenoptera, Apoidea) of the island of Cyprus. ZooKeys. 2020;924:1.

50. Sparrow DJ, John E. An introduction to the wildlife of Cyprus: Terra Cypria; 2016.

51. Christodoulou CS. The impact of Acacia saligna invasion on the autochthonous communities of the Akrotiri salt marshes. University of Central Lancashire, Preston. 2003.

52. Tsintides T, Christodoulou CS, Delipetrou P, Georghiou K. The red data book of the flora of Cyprus. Cyprus Forestry Association Lefkosia 465. 2007.

53. Termaat T, van Strien AJ, van Grunsven RHA, De Knijf G, Bjelke U, Burbach K, et al. Distribution trends of European dragonflies under climate change. Divers Distrib. 2019;25(6):936–50. doi: https://doi.org/10.1111/ddi.12913.

54. Sevilleja C, Collins S, Warren M, Wynhoff I, Van Swaay C, Dennis E, et al. European Butterfly Monitoring Scheme (eBMS): network development. Technical report. Butterfly Conservation Europe and ABLE/eBMS, 2020.

55. Hand R, Hadjikyriakou GN, Christodoulou CS. (continuously updated): Flora of Cyprus – a dynamic checklist Published at http://www.flora-of-cyprus.eu/2011-2020 [cited 2020 09/10/2020]. Available from: http://www.flora-of-cyprus.eu/.

56. Roy HE, Peyton J, Aldridge DC, Bantock T, Blackburn TM, Britton R, et al. Horizon scanning for invasive alien species with the potential to threaten biodiversity in Great Britain. Glob Change Biol. 2014;20(12):3859–71. doi: 10.1111/gcb.12603. PubMed PMID: WOS:000344375700026.

57. Bell S, Marzano M, Cent J, Kobierska H, Podjed D, Vandzinskaite D, et al. What counts? Volunteers and their organisations in the recording and monitoring of biodiversity. Biodivers Conserv. 2008;17(14):3443–54. doi: 10.1007/s10531-008-9357-9.

58. Moreira F, Allsopp N, Esler KJ, Wardell-Johnson G, Ancillotto L, Arianoutsou M, et al. Priority questions for biodiversity conservation in the Mediterranean biome: Heterogeneous perspectives across continents and stakeholders. Conservation Science and Practice. 2019;1(11):e118. doi: https://doi.org/10.1111/csp2.118.

59. August T, Harvey M, Lightfoot P, Kilbey D, Papadopoulos T, Jepson P. Emerging technologies for biological recording. Biological Journal of the Linnean Society. 2015;115(3):731–49.

60. Sutherland WJ, Roy DB, Amano T. An agenda for the future of biological recording for ecological monitoring and citizen science. Biological Journal of the Linnean Society. 2015;115(3):779–84. doi: 10.1111/bij.12576.

61. Giannakopoulos C, Hadjinicolaou P, Kostopoulou E, Varotsos K, Zerefos C. Precipitation and temperature regime over Cyprus as a result of global climate change. Advances in Geosciences. 2010;23:17–24.

62. Hadjinicolaou P, Giannakopoulos C, Zerefos C, Lange MA, Pashiardis S, Lelieveld J. Mid-21st century climate and weather extremes in Cyprus as projected by six regional climate models. Regional Environmental Change. 2011;11(3):441–57.

63. Kleitou P, Crocetta F, Giakoumi S, Giovos I, Hall-Spencer JM, Kalogirou S, et al. Fishery reforms for the management of non-indigenous species. Journal of Environmental Management. 2021;280:111690. doi: https://doi.org/10.1016/j.jenvman.2020.111690.

64. Roy HE, Preston CD, Roy DB. Fifty years of the Biological Records Centre. Biological Journal of the Linnean Society. 2015;115(3):469–74. doi: 10.1111/bij.12575.

65. Franklin J, Serra-Diaz JM, Syphard AD, Regan HM. Big data for forecasting the impacts of global change on plant communities. Glob Ecol Biogeogr. 2017;26(1):6–17. doi: https://doi.org/10.1111/geb.12501.

66. Franz NM, Sterner BW. To increase trust, change the social design behind aggregated biodiversity data. Database. 2018;2018. doi: 10.1093/database/bax100.

67. Hayhow D, Eaton M, Stanbury A, Burns F, Kirby W, Bailey N, et al. State of nature 2019. 2019.

